# Evolution of assortative mating following selective introgression of pigmentation genes between two *Drosophila* species

**DOI:** 10.1101/2022.01.14.476347

**Authors:** Jean R. David, Erina A. Ferreira, Laure Jabaud, David Ogereau, Héloïse Bastide, Amir Yassin

## Abstract

Adaptive introgression is ubiquitous in animals but experimental support for its role in driving speciation remains scarce. In the absence of conscious selection, admixed laboratory strains of *Drosophila* asymmetrically and progressively lose alleles from one parental species and reproductive isolation against the predominant parent ceases after 10 generations. Here, we selectively introgressed during one year light pigmentation genes of *D. santomea* into the genome of its dark sibling *D. yakuba*, and vice versa. We found that the pace of phenotypic change differed between the species and the sexes, and identified through genome sequencing common as well as distinct introgressed loci in each species. Mating assays showed that assortative mating between introgressed flies and both parental species persisted even after four years (∼ 60 generations) from the end of the selection. Those results indicate that selective introgression of as low as 0.5% of the genome can beget morphologically-distinct and reproductively-isolated strains, two prerequisites for the delimitation of new species. Our findings hence represent a significant step towards understanding the genome-wide dynamics of speciation-through-introgression.

## Introduction

In sexually reproducing organisms, speciation begins when extrinsic or intrinsic barriers significantly reduce gene flow between populations and ends with the evolution of pervasive phenotypic differences delimiting the nascent species (Coyne and Orr 2004; The Marie Curie SPECIATION Network 2012; Kulmuni et al. 2020). The pace of this process can be dramatically accelerated if the diagnostic characters also contribute, either directly or through genetic linkage, to reproductive isolation. The search for such traits, which were dubbed ‘magic’, has been a ‘holy grail’ in speciation genetics (Servedio et al. 2011; Smadja and Butlin 2011; Thibert-Plante and Gavrilets 2013; Martin and Richards 2019). However, how such traits form is enigmatic, and theory predicts that substantial degrees of geographical isolation and long times of divergence are necessary for the build-up of genetic barriers to reproduction (Richards et al. 2019). Therefore, it has been argued that adaptive introgression, *i*.*e*. the exchange of beneficial alleles between species with intermediate levels of reproductive isolation (Hedrick 2013), could significantly shorten the time of speciation. Introduced alleles could epistically interact with the host genome leading to the rapid formation of populations that are phenotypically distinct and reproductively isolated from the parental species (Abbott et al. 2013; Schumer et al. 2014; Payseur and Rieseberg 2016; Richards et al. 2019). In spite of the growing evidence for the ubiquity of interspecific gene flow unraveled by recent comparative genomic studies in plants and animals (Lamichhaney et al. 2015; Racimo et al. 2015; Leducq et al. 2016; Pease et al. 2016; Schumer et al. 2018; Edelman et al. 2019), experimental tests for the role of adaptive introgression in the evolution of reproductive barriers are rare. Indeed, two recent reviews on experimental speciation had barely addressed the question of adaptive introgression (Fry 2009; White et al. 2020).

For nearly 100 years, *Drosophila* species have been a primary model for the experimental study of speciation (Mallet 2006; Castillo and Barbash 2017). Introgression between species with incomplete reproductive isolation has long been used to identify the quantitative trait loci (QTL) responsible for phenotypic differences and reproductive barriers (e.g., Tanaka et al. 2015; Ding et al. 2016; Shahandeh and Turner 2020; Massey et al. 2021). In those experiments, two species are crossed and their fertile F_1_ hybrid females are backcrossed to one parental species for one or a few generations. Introgressed genomic regions are then assessed using molecular markers and isogenic lines are produced via inbreeding to test for the statistical association with the phenotype of interest. Such short-term introgression does not inform us much on how introgression can lead to the origin of new species. Indeed, whereas F_1_ hybrid males are sterile, a proportion of males issued from the first backcross are often fertile. When those males are left to mate with the backcross females, the proportion of sterile males progressively diminish each generation. In the absence of conscious selection on a particular introgressed phenotype, alleles from one parent, usually the one that was not used in the backcross, are gradually purged out in less than 20 generations (David et al. 1976; Amlou et al. 1997; Matute et al. 2020). Contrary to those experimental observations, comparative genomics have unraveled strong evidence for genetic introgression between many *Drosophila* species pairs (Lohse et al. 2015; Turissini and Matute 2017; Schrider et al. 2018; Mai et al. 2020), with the traces of introgression sometimes persisting for millions of years (Suvorov et al. 2021).

To test for the effect of adaptive introgression on speciation, one should identify an easily measurable phenotype distinguishing a pair of species, deliberately select it in backcross flies for several generations, and then quantify the degree of reproductive isolation of introgressed flies with both parental species. Unfortunately, most sister *Drosophila* species are usually recognizable only on the basis of subtle differences in their genitalia whose dissection and measuring are quite difficult and laborious (Yassin 2021). A striking exception is the case of the species pair of *D. yakuba* and *D. santomea*, which, in addition to genital differences, also shows a contrasting pigmentation pattern (Lachaise et al. 2000). Both species lack the characteristic sexual dimorphism of pigmentation found in all other species of the *melanogaster* subgroup, where the last abdominal segments of the females are lighter than those of the males. Those segments are equally dark or equally light in both sexes of *D. yakuba* and *D. santomea*, respectively (Figure 1A-D). Both species can mate readily in the laboratory, producing fertile hybrid females but sterile males, and there is strong evidence from field studies and population genomics that hybridization takes place also in the wild on the island of Sao Tomé where *D. santomea* is endemic (Cariou et al. 2001; Llopart et al. 2005, 2014; Turissini and Matute 2017). Leveraging the crossability of the two species, short-term introgression experiments were used to identify the QTL underlying their morphological differences (Coyne et al. 2004; Carbone et al. 2005; Peluffo et al. 2015; Nagy et al. 2018; Liu et al. 2019) and reproductive isolation (Moehring et al. 2006b,a; Cande et al. 2012). Introgressing dark pigmentation alleles of *D. yakuba* in the genome of the lightly pigmented *D. santomea* indicated that at least 5 loci were responsible for the striking pigmentation difference, namely the melanin-synthesis genes *yellow* (*y*), *tan* (*t*) and *ebony* (*e*) and the transcription factors *Abdominal-B* (*Abd*-*B*) and *POU-domain motif 3* (*pdm3*) (Liu et al. 2019). Remarkably, long-term introgression experiments between *D. santomea* and *D. yakuba* showed, that in the absence of conscious selection on any of their morphological differences, reproductive isolation with the parental species may persist for 10 generations (Comeault and Matute 2018), but at generation 20, introgressed flies completely resemble their *D. yakuba* parent with no trace of isolation (Matute et al. 2020).

**Figure 1.**
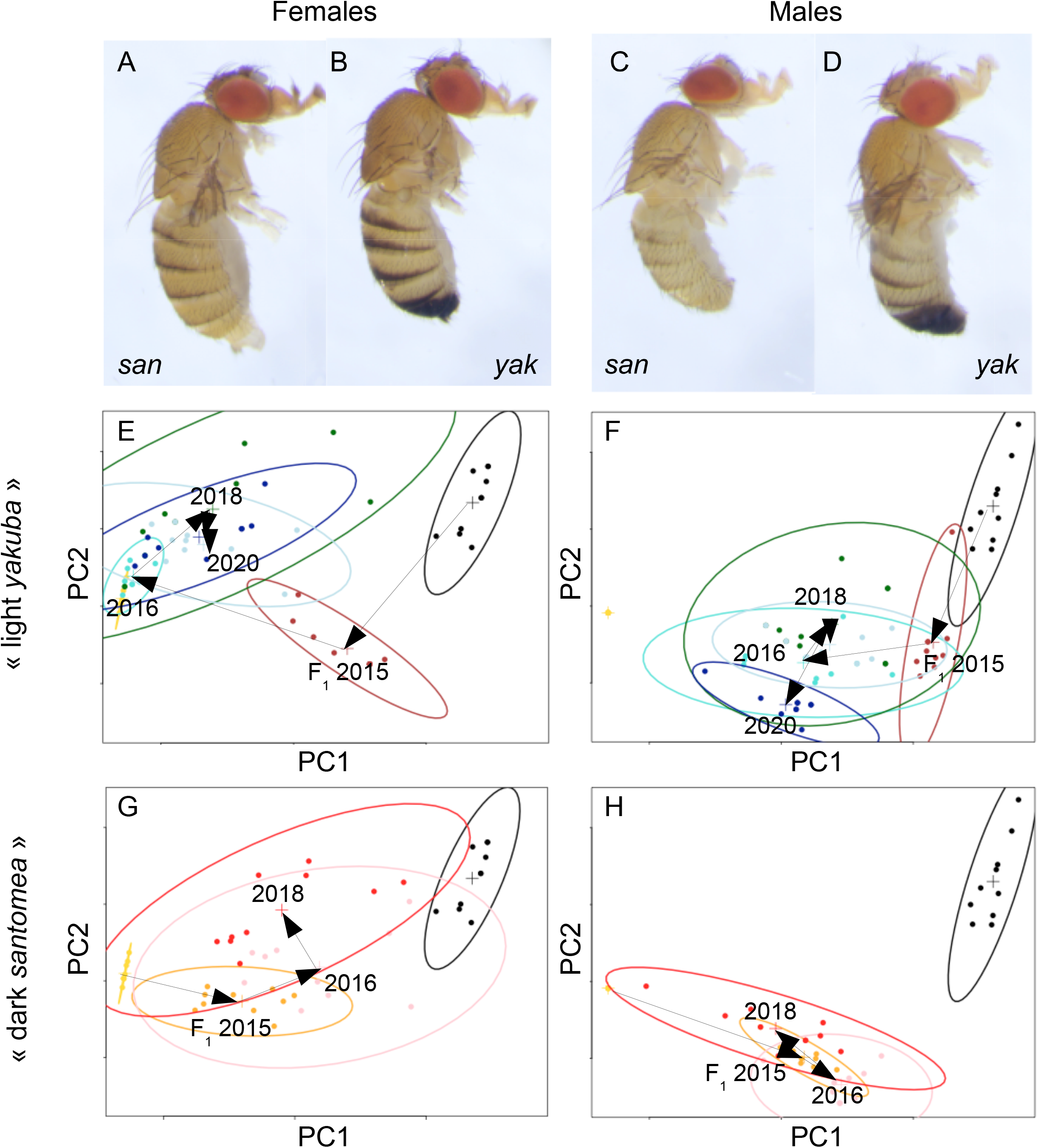
(A-D) Photomicrographs of females and males of the parental species, light *Drosophila santomea* (A,C) and dark *D. yakuba* (B,D). (E-H) Pigmentation introgression trajectories in the “light *yakuba*” (E,F) and the “dark *santomea*” (G,H) experiments. (E-H) Principal Component Analysis (PCA) of pigmentation scores on six successive abdominal segments per individual was conducted on combined males and females data but each sex per experiment was presented in a separate panel according to the coordinates of the two first principal components. In each panel, 95% confidence ellipses for the two parental species are shown in yellow (*D. sanromea*) and black (*D. yakuba*). Colors refer to F_1_ hybrids issued from the cross between female *yakuba* x male *santomea* (brown), *BCyak*^*2016*^ (turquoise), *BCyak*^*2018*^ (dark green), *BCyak*^*selD_2020*^ (dark blue), *BCyak*^*CC_2020*^ (light blue), F_1_ hybrids issued from the cross between female *santomea* x male *yakuba* (orange), *BCsan*^*2016*^ (pink) and *BCsan*^*2018*^ (red). Arrows indicate the trajectory of pigmentation changes in each panel.

In 2015, our late colleague Jean R. David (1931-2021) started two long-term introgression experiments. In the first one, he deliberately introgressed light *D. santomea* alleles in the genome of dark *D. yakuba*, whereas in the second experiment he performed the opposite introgression, *i*.*e*. introgressing dark *D. yakuba* alleles in the genome of light *D. santomea*. In this paper, we report the progress of his 5-year experiments and the results of sequencing two lines from the first experiment. We show through behavioral assays that introgression of as low as 0.5% of the genome has been sufficient to produce flies that were morphologically and behaviorally distinct from both parental species, even after 60 generations from the end of selection. We discuss the relevance of our work to the role of adaptive introgression in speciation.

## Materials and Methods

### Generation of introgression lines

Two experiments were conducted from reciprocal crosses between a strain of *D. yakuba*, which was collected by L. Tsacas from Kounden, Cameroon in 1966, and *D. santomea* from the type laboratory strain collected by D. Lachaise from Sao Tomé Island in 1998. Strains and experimental lines were reared at 21°C on a standard *Drosophila* medium kept in culture bottles at a density of ∼1000 flies.

For the “light *yakuba*” experiment: virgin *D. yakuba* females were crossed to *D. santomea* males. Fertile F_1_ females were mated to *D. yakuba* Kounden males, and the progeny called backcross to *yakuba* (*BCyak*). Backcross flies contained a small proportion (not determined) of fertile males. Those flies were used as a mass population to produce a self-reproducing strain. After a second generation of mass culture, phenotypes were observed on anaesthetized, 3-5 days old flies, and we assumed that most females had already copulated, many of them with fertile males. Selection was made on females only, who were far more variable than males. At each generation ∼50 females with the lightest phenotype were transferred to lay eggs in new culture bottles. Precise phenotypic measurements were not done on regular basis and the progress of selection (if any) was not monitored. However, from our empirical observations, the selection was not efficient; each generation, the light females produced the same proportion of light and dark flies. This result persisted for more than a year (∼ 15 generations). Then, some positive effects were observed: pigmentation of the females became lighter, and also some effects were found on the males, who also could be selected, leading to the establishment of an introgressed *D. yakuba* strain in 2016 (hereafter *BCyak*), quite lighter than the typical *D. yakuba*, especially for the females. However, after two years from the end of selection, female pigmentation slightly increased, attaining the levels of those found in F_1_ hybrids. So a second round of selection on both males and females restarted in 2018, leading to two new derived introgression strains denoted *BCyak*^*CC*^ and *BCyak*^*selD*^ for flies selected for their light and dark abdomen, respectively.

For the “dark *santomea*” experiment: virgin *D. santomea* females were crossed to *D. yakuba* males. The fertile F_1_ females were backcrossed to *D. yakuba* males, and the progeny was reared as a mass culture. Selection started by keeping females with a slightly dark abdomen, but the progress was very slow and took more than a year. Interestingly, the pigmentation of the males increased more rapidly than that of the females, and after about half a year males were also included in selection. In 2016, an introgressed *D. santomea* strain, darker than the typical *D. santomea*, especially for males, was established and denoted *BCsan*.

Throughout the introgression experiments, no samples were archived frozen or in alcohol for genome sequencing and subsequent behavioral assays. Following the perturbations related to the COVID-19 pandemic lockdowns in early 2020, and the deterioration of Jean David’s health later that year, only two strains, denoted BCyak and BCsan were present at the time of genome sequencing in December 2020 and behavioral assays. Those two strains along with those of the parental species were used for genome sequencing and subsequent mapping of introgressed loci. Sequencing unraveled both strains to be predominated by the *D. yakuba* genome, sharing two introgressed *D. santomea* loci at genes known to affect pigmentation (see Results below). Because selection on dark *D. yakuba* alleles in a *D. santomea* background wouldn’t have only fixed light *D. santomea* alleles, we therefore hypothesized that the two strains were derived from the same “light *yakuba*” experiment. This was reconfirmed by checking their male genitalia, which were both of the “*yakuba*” type, in contrast to previous microscopic preparations of *BCsan* strain up to April 2020. A contamination occurring after this date has likely replaced *BCsan* with one of the *BCyak* lines. Because the two strains, *BCyak* and *BCsan*, had two and three fixed *D. santomea* loci (see Results below), the two strains were then denoted *BCyak-2* and *BCyak-3*, respectively.

### Pigmentation scoring and genitalia dissection

Abdominal pigmentation was scored on parental species, reciprocal F_1_ hybrids and the introgression lines following the scoring scheme of David et al. 1990), *i*.*e*. the width of black area at the posterior part of each tergite was visually scored by establishing 11 phenotypic classes from 0 (no black pigment) up to 10 (tergite completely black). Abdominal tergites 2 to 7 as well as tergite 8 (the epigynium) were considered for females and tergites 2 to 6 as well as tergite 9 (the epandrium) were considered for males. For the introgression lines, scoring was made in 2016 at the end of selection, and then once each two years (*i*.*e*. in 2018 and 2020). For each strain, ≥4 days old, 10 females and 10 males were used. Pigmentation scores are provided in Supplementary Table 1. All statistical analyses were conducted using R (R Core Team 2016).

We also aimed to quantify subtle differences in pigmentation intensity between the two strains that were sequenced in 2020, *i*.*e. BCyak-2* and *BCyak-3*. For this, flies were killed in 70% ethanol and wings and legs removed using a pair of forceps. Each fly was then individually placed on its left side in 2 mL 70% ethanol solution in an excavated glass block and photographed under a binocular Leica stereoscope provided with a digital camera connected to a computer. Flies were photographed and grey scale intensity was measured using ImageJ (Abramoff et al. 2004) after manually defining the contour of each abdominal tergite.

The two parental species differ in their male genital traits, with the most easily traceable character being the loss of a pair of hypandrial (sternite 9) bristles in *D. santomea* (Nagy et al. 2018). At the end of selection in 2016, we dissected the male genitalia of the introgression strains and found that the presence or absence of the hypandrial bristles followed the direction of the backcross, *i*.*e*. present in *BCyak* and absent in *BCsan*. Male genitalia were then routinely dissected on a regular basis to guarantee the distinction between the lines of the two experiments.

### Genome sequencing and analysis of two introgressed BCyak strains

For the two strains *BCyak-2* and *BCyak-3*, genomic DNA was extracted from 30 flies using standard DNA extraction kit protocol Nucleobond AXG20 (Macherey Nagel 740544) with NucleoBond Buffer Set IV (Macherey Nagel 740604). DNA was then sequenced on Illumina Novaseq6000 platform (Novogene UK company limited). In order to update the current reference genome of *D. yakuba* v1.05 retrieved from Flybase (https://flybase.org/, Thurmond et al. 2019), we compared this version to a genome of the same *D. yakuba* strain that was sequenced and assembled using hybrid short-read (Illumina) and long-read (Oxford Nanopore) method (http://flyseq.org; Kim et al. 2021). We used assembly-to-assembly command in Minimap2 (Li 2018) to generate a PAF file, based on which we attributed each new ≥ 100 kb-long contig to the corresponding 1.05 chromosomal arm according to the longest homology tract. We also mapped each coding DNA sequence (CDS) to the new contigs using Blast (Altschul et al. 1997) in order to localize previously unmapped 1.05 contigs and genes. For each chromosome, assembled scaffolds were then ordered according to the cytological map of *D. yakuba* in (Lemeunier and Ashburner 1976). This resulted into a newly assembled reference genome of *D. yakuba* (cf. Supplementary Table 2) that we used for mapping introgressed loci.

Minimap2-generated SAM files were converted to BAM format using samtools 1.9 software (Li et al. 2009). The BAM files were then cleaned and sorted using Picard v.2.0.1 (http://broadinstitute.github.io/picard/). We generated synchronized files for the 20 *D. y. yakuba* lines using Popoolation 2. We then used a customized Perl script to extrapolate allele frequencies to 2 diploid counts for each strain, after excluding sites with less than 10 reads and alleles with frequencies less than 25% for the total counts using a customized Perl script (cf. Ferreira et al. 2021). We also excluded tri-allelic sites for each line. We then parsed the parental strains for divergent sites, *i*.*e*. sites with distinct alleles fixed in each strain, and estimated the ancestry proportion at each site in the two introgressed strains in 50 kb-long windows.

### Mating behavioral assays

We estimated precopulatory reproductive isolation between the two parental and the two introgressed strains, *Bcyak-2* and *BCyak-3*, using both no choice and two-choice analyses for both sexes. For no choice analyses, 3-4 days old virgin males and females of all strains were introduced in pairs in individual food vials at around 9:00 AM, and observed for two hours. Mating pairs were counted for each mating pair. For each possible combination of pairs, 20 vials were tested. The proportion of successful matings in intraspecific pairs of *D. yakuba* was considered as the expected proportion, and a chi-squared test comparing the observed proportions of successful mating involving an introgressed and a parental fly for each inter-strain combination.

Two-choice analyses were conducted for both males and females. For a given sex, a virgin fly was introduced into an individual vial along with two virgin flies from the opposite sex, with one being from the same strain as the tested fly and one from another strain. Copulations were observed also for two hours, and once copulation started flies were anesthetized under slight CO_2_, and the identity of the mating and the un-mating flies identified. In some instances, e.g., those involving a *D. santomea* male, no marking was needed. For most other cases, flies were individually left to feed in vials with artificial food blue or red colorants (Sainte Lucie co., France) 24 hours before the start of the experiment as in Comeault and Matute (2018). A chi-squared test was then conducted for each strains combination to test the deviation from parity between homo- and hetero-gamic successful matings.

## Results

### Experimental hybridization led to sexually dimorphic, phenotypically distinct introgression lines

The trajectories of pigmentation evolution during the two 5-year introgression experiments are given in Figure 1E-H in terms of the PCA of pigmentation scores. The first principal component (PC1) explained 75% of the variance. It mostly correlated with the pre-penultimate and penultimate segments (*i*.*e*. segments 6 and 7 in females and 5 and 6 in males) at 0.56 and 0.78, respectively. The second principal component (PC2) explained 13% of the variance, and it mostly correlated with the ultimate segment of the body (*i*.*e*. the female epigynium and the male epandrium) at 0.81. The trajectories differed according to the direction of selection and the sex.

At the end of selection in 2016, introgressed “light *yakuba*” females (Figure 1E) were much lighter than the parental *D. yakuba* (*t*-test for the sum of segments 6 and 7 = 56.65, *P* < 2.2 × 10^−16^). They almost resembled *D. santomea* females, although they were still darker from the later species (*t* = 2.59, *P* = 0.029). Interestingly, all the segments were quite similar, and the last one, *i*.*e*. the epigynium or tergite 8, which is very dark in *D. yakuba* was the lightest in the introgressed females (*t* = 23.24, *P* < 2.4 × 10^−9^). The posterior segments of introgressed males (Figure 1F) were lighter than *D. yakuba* (*t*-test for the sum of segments 5 and 6 = 9.25, *P* < 4.3 × 10^−6^) but still much darker than *D. santomea* (*t* = 10.85, *P* < 1.8 × 10^−6^). However, the last segment, *i*.*e*. the epandrium or tergite 9, became almost completely light (*t* = 10.16, *P* < 1.7 × 10^−6^), as in *D. santomea* (*t* = 1.00, *P* = 0.34). For the “dark *D. santomea*” experiment, introgressed females (Figure 1G) at the end of selection in 2016 were darker than the parental *D. santomea* (*t*-test for the sum of segments 6 and 7 = 10.11, *P* < 3.3 × 10^−6^), but not as dark as *D. yakuba* (*t* = 7.60, *P* < 1.8 × 10^−5^). The males (Figure 1H), on the other hand, had much darker posterior abdomen (*t*-test for the sum of segments 5 and 6 = 21.34, *P* < 5.1 × 10^−9^), yet still lighter than *D. yakuba* (*t* = 10.96, *P* < 4.1 × 10^−7^). The last segments in both sexes were completely light as in *D. santomea*. Remarkably, introgressed females from both experiments significantly differed (*t* = 9.46, *P* < 3.5 × 10^−6^), but not introgressed males (*t* = 1.99, *P* = 0.065).

After two years from the end of selection in 2016, both experiments tended toward pigmentation values of the ancestral backcross parent, but at a much slower rate. This was most pronounced in females of the “light *yakuba*” experiment (*t* = 2.79, *P* = 0.021), but not in males (*t* = 1.02, *P* = 0.321), and in males of the “dark *santomea*” experiment (*t* = 3.42, *P* < 0.004), but not in females (*t* = 1.63, *P* = 0.121). For the second round of selection in the “light *yakuba*” experiment, starting in 2018, the two strains *BCyak*^*CC*^ and *BCyak*^*selD*^ very slightly differed only for male pigmentation of segments 5 and 6 in 2020 (*t* = 2.19, *P* = 0.042). This indicated that selection has attained its limits very rapidly in 2016, but morphological differences between introgressed flies and their parental species persisted for more than 60 generations after selection.

### Two and three D. santomea *loci were fixed in the two light* D. yakuba *strains*

As stated in the Materials and Methods, we sequenced in December 2020 the genome of the two remaining introgressed strains in the laboratory, which were named *BCyak* and *BCsan*. We then estimated the ancestry proportion of both parental species across the genome. This showed that both strains belonged to the “light *yakuba*” experiments, bearing only 5-6% alleles from *D. santomea*. The two strains showed almost the same profile of *D. santomea* introgression tracts, which were classified either as fixed or nearly fixed (*D. santomea* ancestry ≥75%) and intermediate (*D. santomea* ancestry ≥40%) (Table 1; Figure 2). The two strains were called *BCyak-2* and *BCyak-3* in reference to the number of fixed or nearly fixed introgression loci.

**Table 1.** Coordinates according to the *Drosophila yakuba* reference genome v.1.05 of *D. santomea* loci that were fixed (F) or segregate at intermediate frequencies (I) in introgressed light *D. yakuba* strains.

| Locus | Length | *BCyak-2* | *BCyak-3* | No. of genes | Candidate(s) |
| --- | --- | --- | --- | --- | --- |
| X:15,000-226,000 | 211 kb | F | F | 22 | *y* |
| X:17,395,000-17,967,000 | 572 kb | F | F | 49 | *t* |
| 2L:16,511,000-18,064,000 | 1553 kb | I | I | 253 | *pdm3* |
| 3L:3,160,000-4,086,000 | 926 kb | --- | F | 168 | *Gug* |
| 3R:19,079,000-21,169,000 | 2090 kb | --- | I | 304 |  |

**Figure 2.**
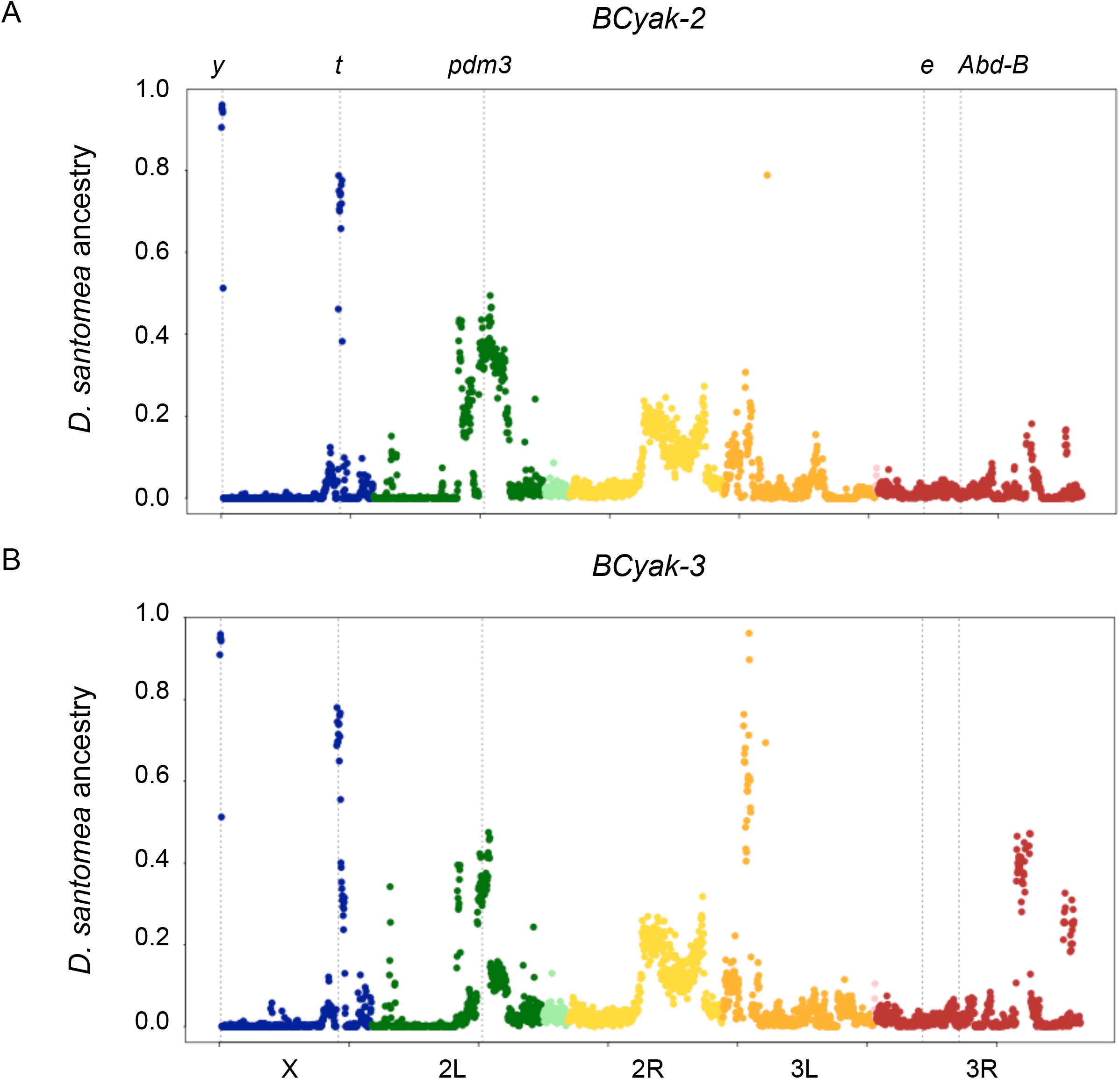
Proportion of *D. santomea* ancestry averaged over 50-kb windows in two introgressed “light *yakuba*” lines (A) *BCyak-2* and (B) *BCyak-3*. Vertical dotted lines refer to the location of the five pigmentation genes that were identified in Liu et al.’s (2019) “dark *santomea*” investigation.

For *BCyak-2* (Figure 2A), the two fixed loci were both X-linked, each centering on one major melanin-synthesis gene, namely *y* and *t*. A third peak with intermediate frequencies was also present on chromosomal arm 2L and it centered on the *pdm3* transcription factor gene. All of those genes, *y, t* and *pdm3*, were found in the opposite experiment by Liu et al. (2019) who introgressed dark *D. yakuba* alleles into *D. santomea*.

The *BCyak-3* strain had exactly the same introgression profile as *BCyak-2, i*.*e*. fixed *y* and *t* loci and intermediate *pdm3* locus (Figure 2B). However, it had also two differences. First, a locus on chromosomal arm 3L had a high proportion of *santomea* alleles and nearly reached fixation. A second locus on chromosomal arm 3R also had high, yet intermediate proportions. None of those two loci harbors any of the previously identified genes known to affect pigmentation differences between *D. santomea* and *D. yakuba* (Liu et al. 2019). However, the 3L locus centered on a transcription factor, *Grunge (Gug)*, which controls the expression of *t* and *e* in *D. melanogaster* (Rogers et al. 2014), and it is therefore a candidate pigmentation locus. There are no candidate pigmentation genes in the 3R locus with intermediate frequency in *BCyak-3*.

The two strains were likely derived from the *BCyak*^*CC*^ and *BCyak*^*selD*^ strains, which corresponded to the second round of selection in the “light *yakuba*” experiment, and which by 2020 slightly differed in male pigmentation (see above). However, the two sequenced strains, *BCyak-2* and *BCyak-3*, did not show significant difference in pigmentation, even when more numerical analyses were used to quantify melanization (Figure 3). Nonetheless, both strains showed significant differences with the two parental species for females’ segment 7 and males’ segment 5, and from a single parent for females’ segment 6 and males’ segment 6, resembling *D. santomea* for the former and *D. yakuba* for the later.

**Figure 3.**
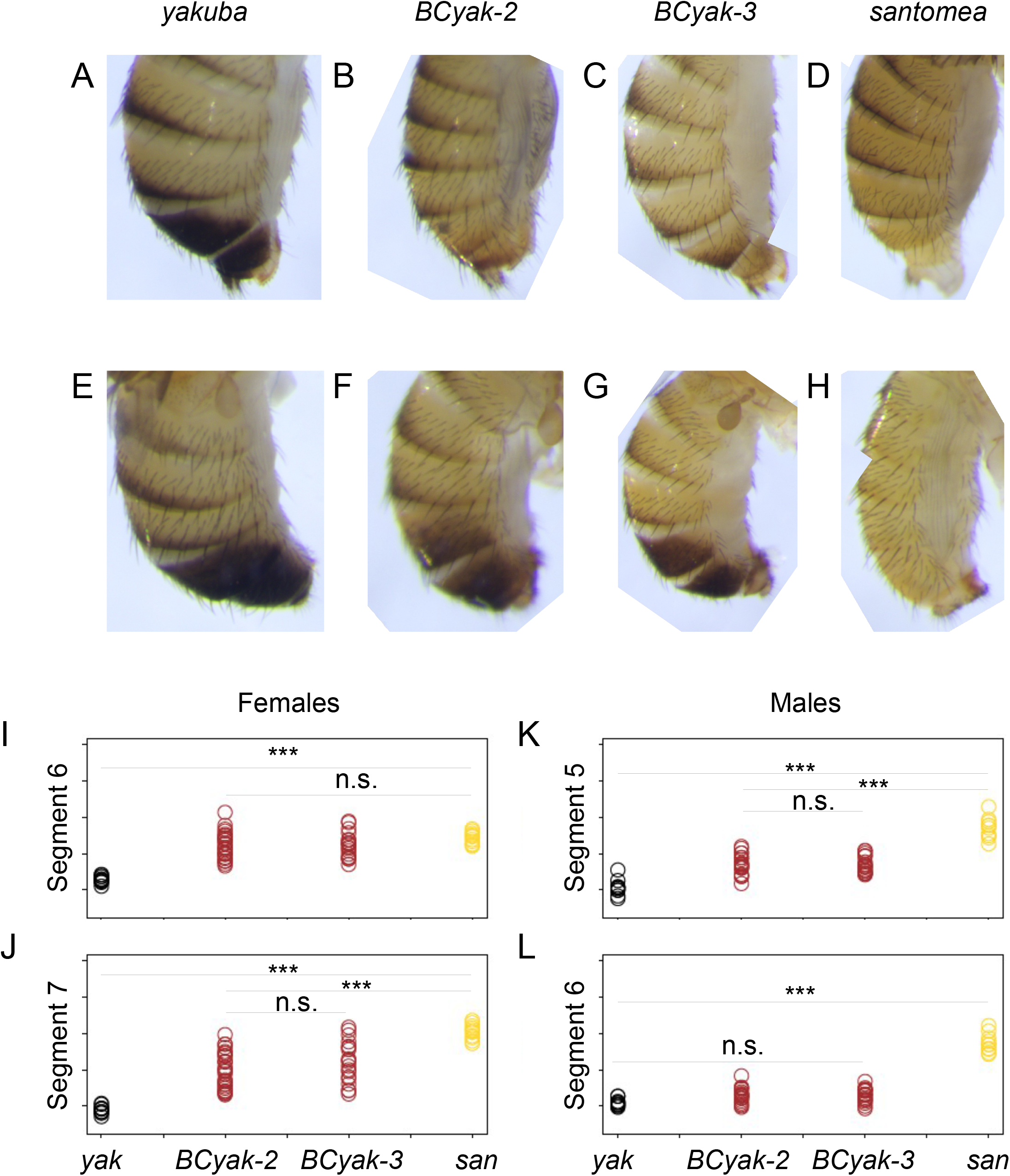
(A-H) Photomicrographs of abdominal pigmentation in males and females of the parental species, *D. yakuba* and *D. santomea*, and the two introgressed “light *yakuba*” lines, *BCyak-2* and *BCyak-3*. (I-L) grayscale intensity of females’ abdominal segments 6 and 7 and males’ abdominal segments 5 and 6. Tukey’s HSD significance level: * < 0.05, ** < 0.01 and *** < 0.001.

### Assortative mating between introgressed strains and parental species

In no-choice experiments, homogamic mating occurred with almost the same frequency between pairs belonging to the same strain/species (70-85%) (Table 2). The two introgressed *yakuba* lines, *BCyak-2* and *BCyak-3*, readily mated with each other. However, a significant low mating success was observed in the cross between *D. yakuba* females and *BCyak-3* males. Interspecific crosses between *D. santomea* and *D. yakuba*, as well as between *D. santomea* females and males from both introgressed lines were significantly low. Remarkably, more successful heterogamic matings were observed in cases involving *D. santomea* males and females from the introgressed *yakuba* lines who have lighter abdomen compared to *D. yakuba*.

**Table 2.**
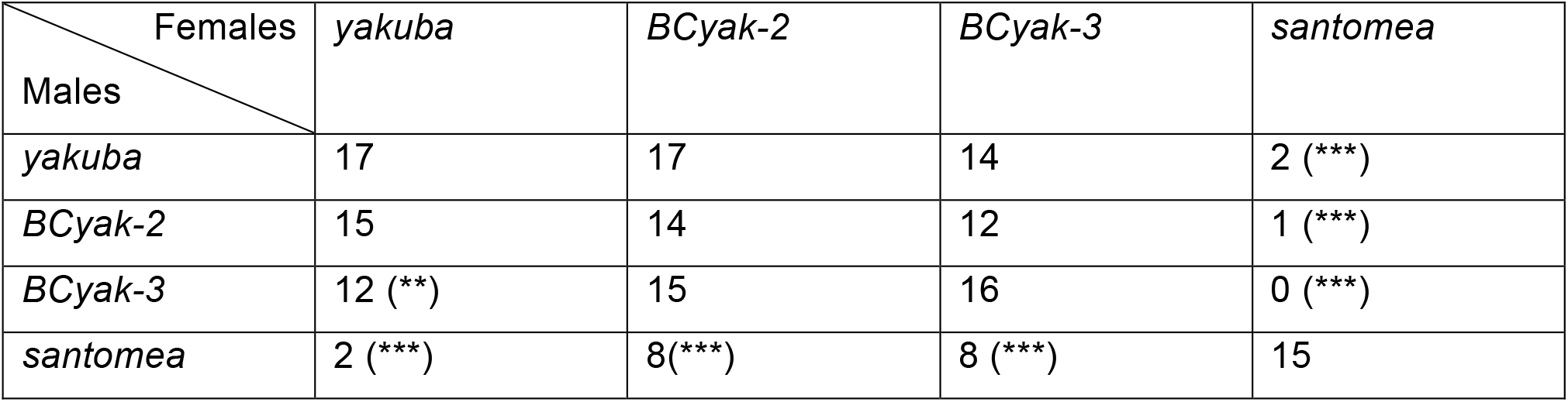
No choice experiment within and between pure parental species, *D. yakuba* and *D. santomea*, and two introgressed “light *yakuba*” strains. 20 copulating pairs were tested per cross. For heterogamic crosses, significant deviation from the homogamic *D. yakuba* cross cross, *i*.*e*. 17 successful crosses out of 20, was estimated using chi-squared test: * < 0.05, ** < 0.01 and *** < 0.001.

For choice experiments, all crosses involving *D. yakuba* and the introgressed lines on the one hand and *D. santomea* on the other hand were significantly homogamic, regardless to the tested sex (Table 3). However, sex-dependent assortative mating was found for all crosses between *D. yakuba* and introgressed strains. In all those crosses, females always showed a higher preference for homogamic males, whereas no significant departure from parity was observed for males.

**Table 3.**
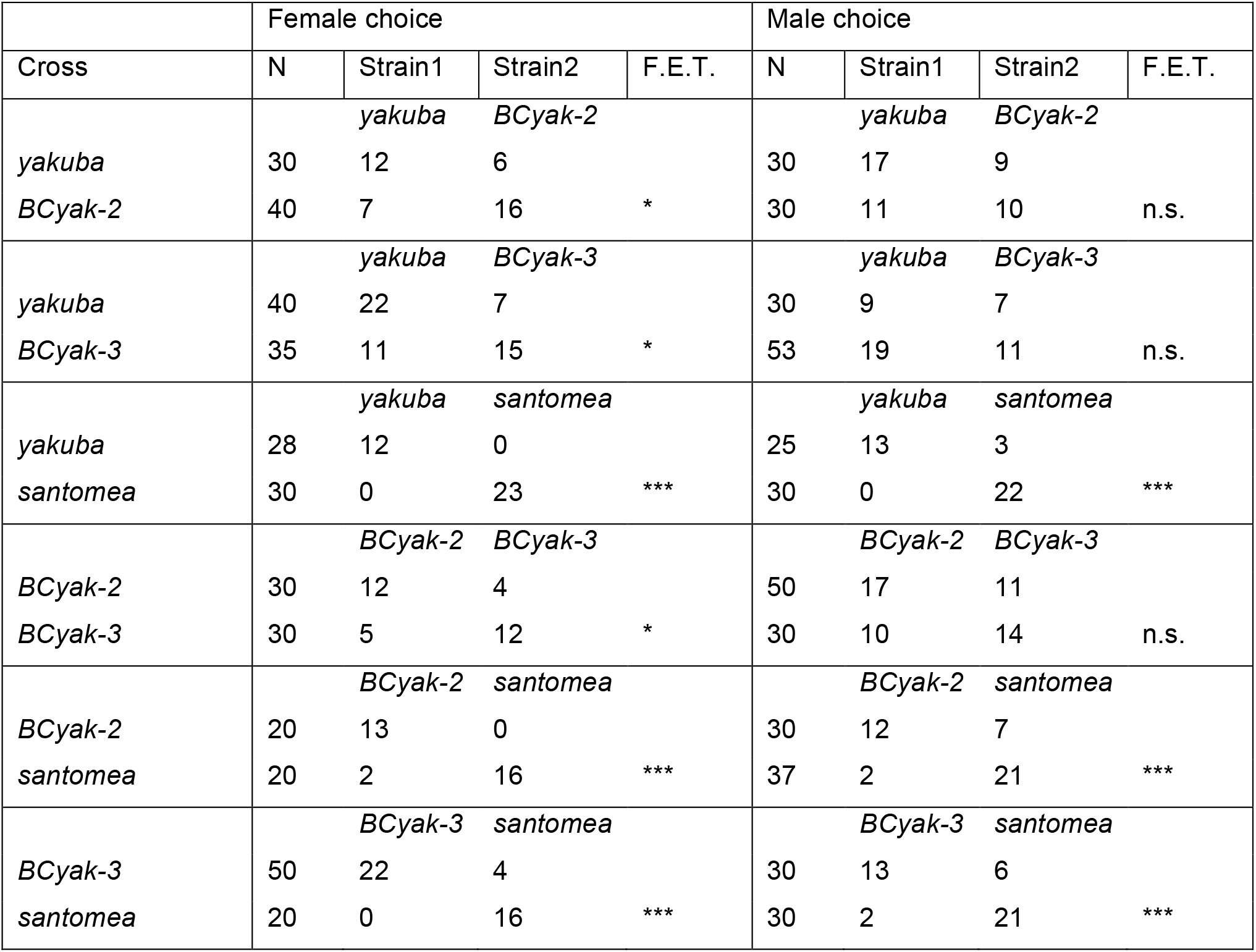
Two-choice mating preference experiments. F.E.T. = significance level of Fisher’s exact test for homogamy in each possible combination: * < 0.05, ** < 0.01 and *** < 0.001.

## Discussion

We reported here the results of five-years experiments to reciprocally introgress genes causing morphological difference between a pair of sister species with a major difference in body pigmentation, and a strong, yet incomplete reproductive isolation. We showed that such introgression was possible and that the limits of selection was attained within only a single year (∼15 generations), with the new phenotypes of the introgressed flies remaining distinct from the parental species. Remarkably and contrary to previous studies with no conscious selection on a morphological trait (David et al. 1976; Amlou et al. 1997; Matute et al. 2020), assortative mating persisted in the introgressed flies even after four years from the end of selection (∼60 generations).

The success of selective introgression might strongly depend on the nature of the phenotype. Pigmentation can easily be scored and measured and its variation often has a simple, oligogenic architecture (Massey and Wittkopp 2016). By contrast, when Amlou et al. (1997) tried to introgress resistance to a fruit toxin from *D. sechellia* into *D. simulans*, their attempt has failed, likely due to the difficulty of measuring toxicity and to the polygenic nature of survival as a phenotype. Indeed, many known cases of cross-species adaptive introgression involve color variation, e.g., coat in wolves (Anderson et al. 2009), skin and hair colors in humans (Dannemann and Kelso 2017), wing patterns in mimetic butterflies (Edelman et al. 2019), winter-coats in hares (Giska et al. 2019), plumage in pigeons (Vickrey et al. 2018) and wagtails (Semenov et al. 2021), and beaks in Darwin’s finches (Enbody et al. 2021).

Introgressed flies differed from their parents in both the degree of pigmentation but also in resuscitating ancestral sexual dimorphism that was independently lost in the parental species. In most species of the *melanogaster* species subgroup, including the closely-related *D. teissieiri*, males have darker abdomen (Yassin et al. 2021). The loss of sexual dimorphism in *D. santomea* and *D. yakuba* has likely involved different sex-specific regulatory changes affecting similar sets of melanin-synthesis genes. We were not able to sequence our introgressed “dark *santomea*” flies which were lost by mid-2020, but fortunately Liu et al. (2019) have conducted similar experiment and identified at least five genes whose *D. yakuba* alleles darken *D. santomea* male pigmentation. Our introgressed loci in the “light *yakuba*” flies overlapped with three out of these genes, namely the X-linked melanin-synthesis genes *y* and *t* and the autosomal transcription factor *pdm3*. By contrast, we did not detect signal of introgression on either the melanin-synthesis gene *e* or the homeotic transcription factor *Abd-B*, which were identified in “dark *santomea*” (Liu et al. 2019). This was in agreement with Liu et al.’s (2019) observations. *Abd-B*, which has lower expression in *D. santomea*, does not affect *D. santomea* pigmentation genes due to *cis*-regulatory mutations of its melanin-synthesis genes. Similarly, whereas *D. santomea e* has a higher expression associated with the insertion of a helitron in its regulatory sequence, the presence of the same *D. santomea* haplotype in *D. yakuba* does not affect its pigmentation (Liu et al. 2019).

The most intriguing result was the autosomal locus that was fixed or nearly fixed in only one of the two introgressed *BCyak* strains, and which was not identified by Liu et al. (2019) in their “dark *santomea*” flies. This locus contained the transcription factor *Gug*, which may have the opposite effect of *pdm3* on pigmentation intensity and sexual dimorphism. RNA interference (RNAi) silencing of this gene in the abdomen of *D. melanogaster* reduces pigmentation, with the reduction being more pronounced in males, whereas RNAi of *pdm3* increases pigmentation, with the increase being more pronounced in females (Rogers et al. 2014). Whereas *pdm3* is a suppressor of *y* in *D. santomea* (Liu et al. 2019), *Gug* is an enhancer of *t* and a suppressor of *e* in *D. melanogaster* (Rogers et al. 2014). Therefore, it is possible that the gain of female-specific pigmentation in *D. yakuba* was partly due to a down-regulation of *pdm3* whereas the loss of male-specific pigmentation in *D. santomea* was partly due to an up-regulation of *Gug*. The lack of significant difference in pigmentation between *BCyak-2* and *BCyak-3* argues against any role of the 3L locus, including *Gug*, on pigmentation. However, we note that pigmentation analysis of those two strains has been made in December 2021 after at least 18 months from the end of the second round of selection in the “light *yakuba*” experiment. Laboratory experiments and population analyses in *Drosophila* have suggested that balancing selection may act on pigmentation genes, hence restoring their alleles to intermediate frequencies when selection ends (L’Héritier and Teissier 1937; Kalmus 1945; Rendel 1951). For example, pigmentation polymorphism in *D. kikkawai*, which is controlled by the *pdm3* locus (Yassin et al. 2016b), is maintained by heterozygous advantage in experimental populations (Freire-Maia 1964). Similarly, ancient balancing selection on *t* was demonstrated in *D. erecta* (Yassin et al. 2016a). Further isolation from *pdm3* and *t* of the introgressed locus on 3L and subsequent molecular dissection are therefore needed to understand its potential role in pigmentation evolution.

Color-based assortative mating could lead to the loss of sexual dimorphism and ultimately pre-copulatory reproductive isolation. Our results showed that fixation of as low as 0.8 Mb (∼0.5% of the genome) during selection on pigmentation loci has altered mating propensities between pure and introgressed flies. The demonstration of color-based (dis)assortative mating in *Drosophila* has long been problematic (Kopp et al. 2000; Llopart et al. 2002). Our behavioral assays support the presence of color-based assortative mating between *D. yakuba* and *D. santomea*, but in a way that was asymmetric between the sexes and dependent on the degree of divergence. On the one hand, light male *D. santomea* had almost 5-fold success in mating with introgressed light *D. yakuba* females than with dark pure *D. yakuba* in no choice experiments. On the other hand, light females from both introgressed *BCyak-2* and *BCyak-3* showed preference for their own light males over pure dark *D. yakuba* males. This suggests that the two X-linked *y* and *t* loci that were fixed in both strains probably play a role in color-based assortative mating. However, female-limited assortative mating also existed between the introgressed strains *BCyak-2* and *BCyak-3*, in spite of their great coloration resemblance. The fixed autosomal locus in *BCyak-3* may therefore also contain elements affecting behavior. In addition to its possible effect on pigmentation, the transcription factor *Gug* also interacts with another transcription factor, *hairy (h*), which is also located in the same fixed locus, in affecting the size of male genital organs that are used to grasp the females during mating, namely the surstyli (claspers) (Hagen et al. 2021). The effect of pigmentation genes on mating behavior can be attained either directly through pleiotropy or indirectly genetic linkage to other mating phenotypes (Wellenreuther et al. 2014). Pleiotropy should drive more pervasive associations between pigmentation and mating behavior than linkage. A possible source of genetic linkage could have been the physical proximity in the low recombining subtelomeric region of the X chromosome between *y* and the enhancer of *scute (sc)* which led to the loss of the hypandrial bristles and gain of extranumerary sex comb teeth in *D. santomea* males (Nagy et al. 2018). Both characters may be involved in copulation and consequently contribute to mating success or choice. However, we found that this strong linkage was broken during the first year of the selection experiment, dissociating both traits.

In conclusion, our result demonstrate that selective introgression on a morphological phenotype could rapidly lead to the evolution of pervasive behavioral isolation. They hence complement previous *Drosophila* experimental speciation studies, which showed that adaptation from standing variation to contrasting environments could lead the evolution of reproductive isolation (Fry 2009). Pigmentation also responds to diverse natural selection pressures (Bastide et al. 2014) including those that discriminate the ecological niches of *D. santomea* and *D. yakuba* such as temperature, desiccation and UV intensity (Matute et al. 2009; Matute and Harris 2013; Comeault and Matute 2021). Further experimental manipulations, e.g., testing competition between pure and introgressed flies in different environments, coupled with the investigation of post-copulatory isolation barriers, will definitively shed more light on genome dynamics of homoploid speciation in animals, hence bridging experimental studies with empirical field observations in a primary model.

## Conflicts of interest

We declare no conflicts of interest.

## Acknowledgments

This work was partly funded by the French Agence Nationale de la Recherche (ANR) grant number ANR-18-CE02-0008 to A.Y.

## Notes

### Competing Interest Statement

The authors have declared no competing interest.

